# Mindfulness training impacts brain network dynamics linked to stress response in young adolescents

**DOI:** 10.1101/2024.10.16.616959

**Authors:** Julian Gaviria, Zeynep Celen, Mariana Magnus Smith, Lucas Peek, Soraya Brosset, Patrik Vuilleumier, Dimitri Van De Ville, Arnaud Merglen, Paul Klauser, Camille Piguet

**Affiliations:** Department of Psychiatry, Amsterdam UMC; Amsterdam Neuroscience, Amsterdam UMC; Vrije Universiteit, Amsterdam. The Netherlands; Department of Psychiatry; Laboratory for Behavioral Neurology and Imaging of Cognition, department of Basic Neurosciences; department of radiology and medical informatics, University of Geneva. Switzerland; Medical Image Processing Lab, Neuro-X Institute, Ecole polytechnique fédérale de Lausanne (EPFL). Switzerland; Center for psychiatric neuroscience, Department of psychiatry, Lausanne university hospital and University of Lausanne, Lausanne, Switzerland; Division of child and adolescent psychiatry, Department of psychiatry, Lausanne university hospital and University of Lausanne, Lausanne, Switzerland

**Author notes:** **Correspondence** Julian Gaviria.

**Keywords:** Brain networks, mindfulness, dynamic functional connectivity (dFC), depression, stress, adolescents, anxiety

## Abstract

Mindfulness-based interventions (MBI) may lead to lower levels of psychological distress, including depression, anxiety, and stress in adolescents. Past research has advanced the discovery of neural architecture recruited by MBI. However, the brain mechanisms through which mindfulness exerts more resilient responses to social stressors in teens remain unclear. Here, we examined how MBI modulates changes in brain network dynamics following social stress with different affective valence (i.e., neutral, negative, and positive). For this aim, we carried out a longitudinal randomized controlled trial in which non-clinical adolescents underwent MBI for 8 weeks. They completed a psychosocial stress task before and following MBI. Functional magnetic resonance imaging (fMRI) and self-reported measurements of psychological distress were collected in both measurement points (i.e., “pre” and “post” MBI). We computed co-activation patterns on fMRI data to characterize dynamic functional connectivity within whole-brain networks. The results depicted how MBI modulates transient co-activation changes in dorsal medial regions of the brain default network (DN) following the experience of stress. However, these brain changes were not specific to the affective valence of stressful stimuli. The relationship between the DN dynamics and the measurements of psychological distress was mediated by MBI. Globally, our findings support a model in which MBI causally mediate brain-behavior interactions related to psychosocial stress in adolescents.

## Introduction

The present study investigated the impact of mindfulness on brain functional systems underlying psychosocial stress response in adolescence, a critical stage for the onset of multiple mental disorders (1). One out of five adolescents experience mental disorders, a number rapidly increasing in Western societies (2). This vulnerability stems partly from exposing the immature brain to highly stressful situations in adolescence, along with widespread delays in treatment. Adolescence is, therefore, a critical period for mental health interventions (3,4). Mindfulness-based intervention (MBI) is a low-cost intervention effectively used to reduce stress (5–7). Besides being a skill that can be acquired through training, mindfulness is also conceptualized as a personality trait (8,9) which is subject to improvement through continuous mindfulness training (10). Studies on the impact of Mindfulness-Based Interventions (MBI) on adolescents have shown a correlation between mindfulness traits and reduced anxiety, depression, somatic symptoms, improved self-esteem, and sleep quality (11). However, there is limited research on the brain mechanisms involved in this relationship (12). Previous studies have examined brain static functional connectivity (FC) to identify the neural architecture associated with mindfulness. They found positive associations between mindfulness trait and regions in the default network (DN), such as the inferior frontal and orbitofrontal areas (13), as well as negative associations with the thalamus (14) and the precuneus (15).

Recent works investigated whether changes in the dynamics of FC [i.e., dFC (16,17)] captured the effects of mindfulness on the brain. Unlike standard methods, the dFC approach evaluates changes in functional brain patterns (i.e., networks) over time. It is also known for unveiling fine-grained signatures of brain functioning related to affective states in health (18,19) and mood disorders (20,21). Two studies reported a reduction in dFC linked to mindfulness trait: Marusak et al. (22) identified an association between decreased dFC in the salience (SN), the central executive brain network (CEN), and mindfulness trait in children and adolescents. Teng and colleagues (23) found that reduced dFC duration in SN, CEN, and DN areas was linked to greater mindfulness trait and attenuated affective and neuroendocrine stress responses in young adults. These findings paved the way for exploring the impact of mindfulness on the brain’s dFC in youth. However, it remains unclear how the adolescent brain reacts to various types of psychosocial stressors. To investigate these issues, we conducted functional magnetic resonance imaging (fMRI) while adolescents were exposed to acute psychosocial stress, with recovery periods between stressors. We experimentally induced neutral, negative, and positive social stressors using a modified Montreal Imaging Stress Task [MIST(24,25)]. This approach enabled us to explore how sustained carry-over effects of different stressors could impact neural activity and clinical measurements collected after stress, both before and after mindfulness training.

We implemented a recently developed dFC method to obtain quantitative temporal parameters, as well as anatomical characteristics across the whole brain. Specifically, we leveraged a seed-based co-activation patterns approach [CAPs (26,27)], with the bilateral mid-frontal (MFG), inferior frontal gyrus (IFG), and posterior cingulate cortices (PCC) as the brain regions of interest (i.e., ROIs. **Figure 1E**). These regions have been previously related to acute stress response in adolescents (28). The CAPs approach permitted us to track dynamic fluctuations of network activity over time with voxel-wise resolution. We then probed relationships between the resulting CAPs and clinical measures related to depression and anxiety, as well as the causal mediation of mindfulness trait on this brain-behavior relationship to test two complementary hypotheses: (i) the brain functional connectivity exhibits specific patterns (i.e., CAPs) related to mindfulness trait (**Figure 1A**). (ii) These CAPs describe the effects of different types of psychosocial stress (i.e., neutral, negative, and positive) on the brain. In line with past research, we expected that the mindfulness intervention (MBI) induces changes in distributed brain circuits. They include the default network (DN), typically expressed in the resting state, as well as the SN and CEN networks, typically recruited by psychosocial stress (28). More critically, we predicted a significant mediation of mindfulness trait on the relationship between the brain CAPs and clinical scores related to psychological distress.

**Figure 1.**
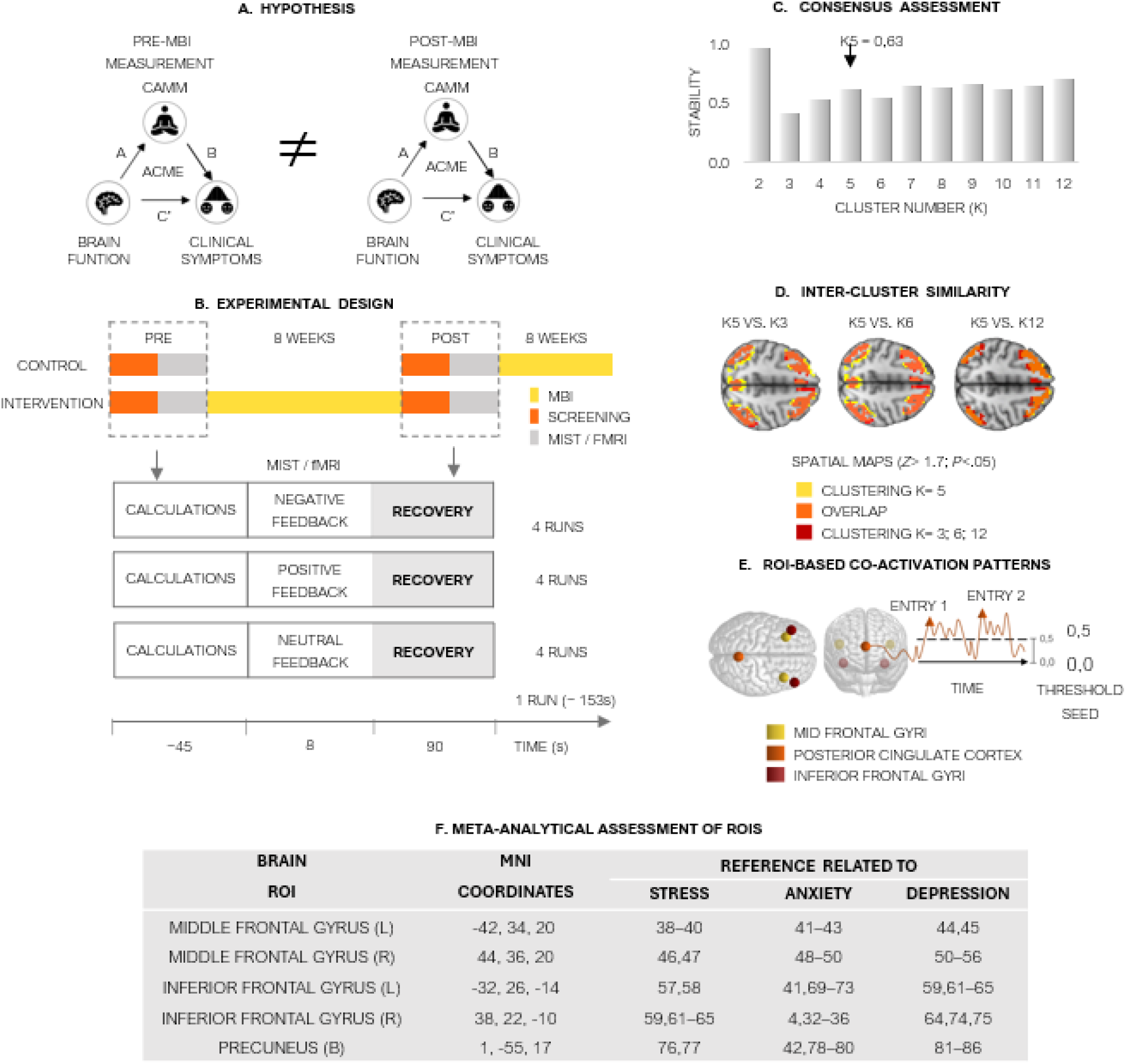
**A**. First research hypothesis: Mindfulness training impacts the relationship between brain function and clinical symptoms in adolescents. This influence is observed through changes (“pre “vs. “post” measurements) in the average causal mediation effect (ACME) of trait mindfulness (CAMM) on the mentioned brain-behavior association. **B**. Experimental design. The longitudinal study with a nested randomized controlled trial included two groups. The “INTERVENTION” group experienced a mindfulness-based intervention (MBI) between the first visit (“PRE” MBI) and the second visit (“POST” MBI). The “CONTROL” group underwent the MBI following the second visit. FMRI and clinical phenotypes were collected in both visits. In addition, the participants performed the Montreal Imaging Stress Task (MIST), which consisted of arithmetic calculations (max. 45 seconds), feedback about the task performance (8 seconds), and lastly, a resting state period (“RECOVERY” (90s)). Three types of feedback were presented: neutral, positive, and negative. **C**. Intercluster similarity analysis. Spatial maps were compared to identify partitions (i.e., CAPs) with the greatest consistency and reliability across different clustering scenarios. E.g., K=5 vs. K=3, K=6, K=12, respectively. **D**. Consensus assessment. The stability plot indicates the clustering quality at the assessed k (from k=2 to k=12). These results provide a quantitative metric suggesting that k=5 is the optimal number of clusters for classifying our fMRI dataset. **E**. ROI-based co-activation patterns. 6-mm spherical brain regions of interest (ROIs) were selected for the CAP analysis: Bilateral mid frontal gyri (yellow. MNI coordinates: -42, 34, 20/44, 36, 20), bilateral posterior cingulate gyr (orange. MNI coordinates: 1, -55, 17), and inferior frontal gyri (red. MNI: 32, 26, -14/38, 22, -10) The activity time-course from each seed was computed separately for each subject, and the frames for which it exceeded a positive threshold (Threshold seed = > 0.5) were tagged. The entry rates quantified their temporal expression over the fMRI scanning. E.g., one CAP derived from the PCC is represented with orange color. This CAP “entered” twice (orange triangles) into one brain state, encompassing the connectivity between the PCC ROI and the whole brain. Namely, it crossed a predetermined threshold twice during the scanning. Further description is provided in (27). **F**. Metanalytical assessment of the predefined ROIs, previously associated with the neural processing of stress, anxiety, and depressive symptoms. The reference numbers correspond to the bibliographical index for each study.

## Methods

### Participants

Seventy French-speaking participants were recruited through a large communication strategy, including social media, adolescent consultation at the hospital, pediatricians, schools, and a website as described in the protocol paper (25). Inclusion criteria: Age (between 13 and 15 years old), no experience in the practice of meditation, no history of chronic somatic or psychiatric illnesses (except for mood disorders resolved for at least 6 months or current anxiety disorder without comorbidities), no psychological follow-up, as well as the absence of contraindications for the various examinations, e.g., neuroimaging. Most of the sample had no lifetime psychiatric diagnosis, while 13 reported mental health difficulties such as past depressive episodes or a current anxiety disorder on the psychiatric diagnostic interview [K-SADS-PL (29)]. The participants had no treatment, and no co-intervention was allowed, see **Table 1**. Five participants (four in the intervention group) were excluded from the final sample due to excessive movement during the neuroimaging recordings.

**Table 1.** Descriptive of the Mindfulteen Study sample. Abbreviations: MBI: Mindfulness-based intervention. GDA: generalized anxiety disorder. COVID: Coronavirus disease 2019. STAI: State/trait anxiety inventory. BDI: Beck depression inventory. DASS: Depression Anxiety Stress Scales. CAMM: Child and Adolescent Mindfulness Measure.

| VARIABLE | Mean (SE) |  |  |  |
| --- | --- | --- | --- | --- |
|  | All (70) | Treatments (33) | Controls (36) | Range |
| Age (years) | 14,30 (0,11) | 14,29 (0,15) | 14,30 (0,15) | 13,07 -16,01 |
| Duration between scans | 99.81 (31.53) | 108.56 (28.95) | 92.03 (32.06) | 49-196 |
| Number of MBI Sessions | 7,49 (0,10) | 7,48 (0,15) | 7,50 (0,13) | 5-8 |
| Gender (F/M) | 41/29 | 22/11 | 18/18 | - |
| Puberty (No/Yes) | 20/47 | 6/27 | 10/26 | - |
| Difficulties (KSADS) (No/Yes) | 56/13 | 29/4 | 27/9 | - |
| Social phobia (No/Yes) | 63/7 | 28/5 | 34/2 | - |
| GAD (No/Yes) | 69/9 | 27/6 | 33/3 | - |
| STAI-S | 36.46 (0.93) | 37.51 (1.49) | 35.56 (1.16) | - |
| STAI-T | 31.77 (0.71) | 31.34 (0.97) | 32.14 (1.02) | - |
| BDI | 9.91 (0.95) | 10.88 (1.61) | 9.09 (1.11) | - |
| DASS-A | 3.52 (0.44) | 3.55 (0.57) | 3.49 (0.67) | - |
| DASS-D | 4.54 (0.52) | 4.74 (0.91) | 4.36 (0.58) | - |
| DASS-S | 5.30 (0.53) | 5.51 (0.86) | 5.11 (0.67) | - |
| CAMM | 27.02 (0.94) | 26.6 (1.4) | 27.4 (1.2) | - |

### Paradigm design

Mindfulteen is a longitudinal cohort study with a nested randomized controlled trial (30). After inclusion, participants were electronically randomized using the sealedenvelope® platform between the “intervention” group and the “control” group (**Figure 1B**). Participants allocated to the control group were engaged in MBI after the waiting period. Assessments were performed before intervention (i.e., “pre-MBI”) and immediately after the intervention (i.e., “post-MBI”) or after the waiting period for the control group, as detailed in **Figure 1B**. The MBI was specifically designed in-house for a young adolescent audience – adapted from standardized meditation programs such as mindfulness-based cognitive therapy [i.e., MBCT (31)] and mindfulness-based stress reduction [i.e., MBSR, (32)]. The program for this intervention consisted of one 90-minute group practice session per week for 8 consecutive weeks (see (30) for detailed program content). Participants were also encouraged to practice individually each day using an in-house smartphone app.

### FMRI data

Our study focused on dynamic functional connectivity (dFC) of the brain computed from resting periods of functional magnetic resonance scanning (fMRI). It was acquired with a 3T Magnetom TIM Trio scanner (Siemens, Germany) at the Brain and Behavior Laboratory at the Faculty of Medicine of the University of Geneva (Switzerland). See SI “Supplementary Methods” section for further description of both data acquisition and preprocessing).

### Task and measures

The fMRI task was an age-adapted version of the modified Montreal Imaging Stress Task (MIST) used previously by our group (24,25). The MIST is a psychosocial stress task with an acute stressor and social evaluative feedback with recovery periods in between conditions. See further description in the SI “Supplementary Methods” section. Each participant was evaluated with the Kiddie schedule for affective disorders and schizophrenia. Present and lifetime version [i.e., K-SADS-PL (33)]. Among self-questionnaires which were administered to participants at each visit: anxiety was assessed by the State-Trait Anxiety Inventory for Children [i.e., STAI-C (34)], depression was assessed by the Beck Depression Inventory [i.e., BDI (35)] and stress was assessed by the Depression and Anxiety Stress Scale [i.e., DASS-21 (36)]; we also used The Strengths and Difficulties Questionnaire to assess global functioning (37). Trait mindfulness was measured with the child and adolescent mindfulness measure [i.e., CAMM (38)].

### Dynamic functional connectivity (dFC)

We first computed functional seed-based co-activation patterns (i.e., CAPs) on resting state fMRI data. We used the CAPs approach because, unlike standard methods, it captures changes in functional brain patterns (i.e., networks) over time. Also, the brain topography depicted by CAPs is data-driven. Namely, the CAPs are not constrained by an aprioristic brain compartmentalization (i.e., functional atlas). Further description of the CAPs approach is provided in **Figure 1** and (26,27). Our brain regions of interest (i.e., ROIs) included the mid-frontal (MFG. MNI coordinates: - 42, 34, 20/44, 36, 20), inferior frontal gyrus (IFG. MNI: 32, 26, -14/38, 22, -10), and posterior cingulate cortices (PCC. MNI: 1, -55, 17). **Figure 1E**. These regions were identified by Corr et al., as highly responsive to acute stress in adolescents (28). In addition, they constitute key hubs of brain networks active at rest, particularly in the default network (DN), salience (SN), and the central executive network (CEN), and have been previously associated with the neural processing of stress, anxiety and depressive symptoms (**Figure 1F**). (39–41) (42–44) (45,46) (47,48) (49–51)(51–57) (58,59)(49,60,61) (60,62–66) (41,67–69) (42,70–74) (65,65,75,76)(77,78) (43,79–81) (82–87)

### Spatiotemporal CAPs generation

One CAP consists of a functional map (i.e., brain network. **Figure 2A**) and one or multiple indicators of temporal variability of this functional network. Here, the “entries” metric quantifies the temporal expression of each brain CAP across different groups and measurements (e.g., **Figure 2B**). “Entry rates” are the number of times the brain enters a specific state of co-activation (**Figure 1E**). Multilevel linear mixed models (LMMs) assessed the temporal expression of our CAPs in terms of the statistical variance of their entry rates. Further description of our LMM modeling strategy is provided in the SI “Supplementary Methods” section.

**Figure 2.**
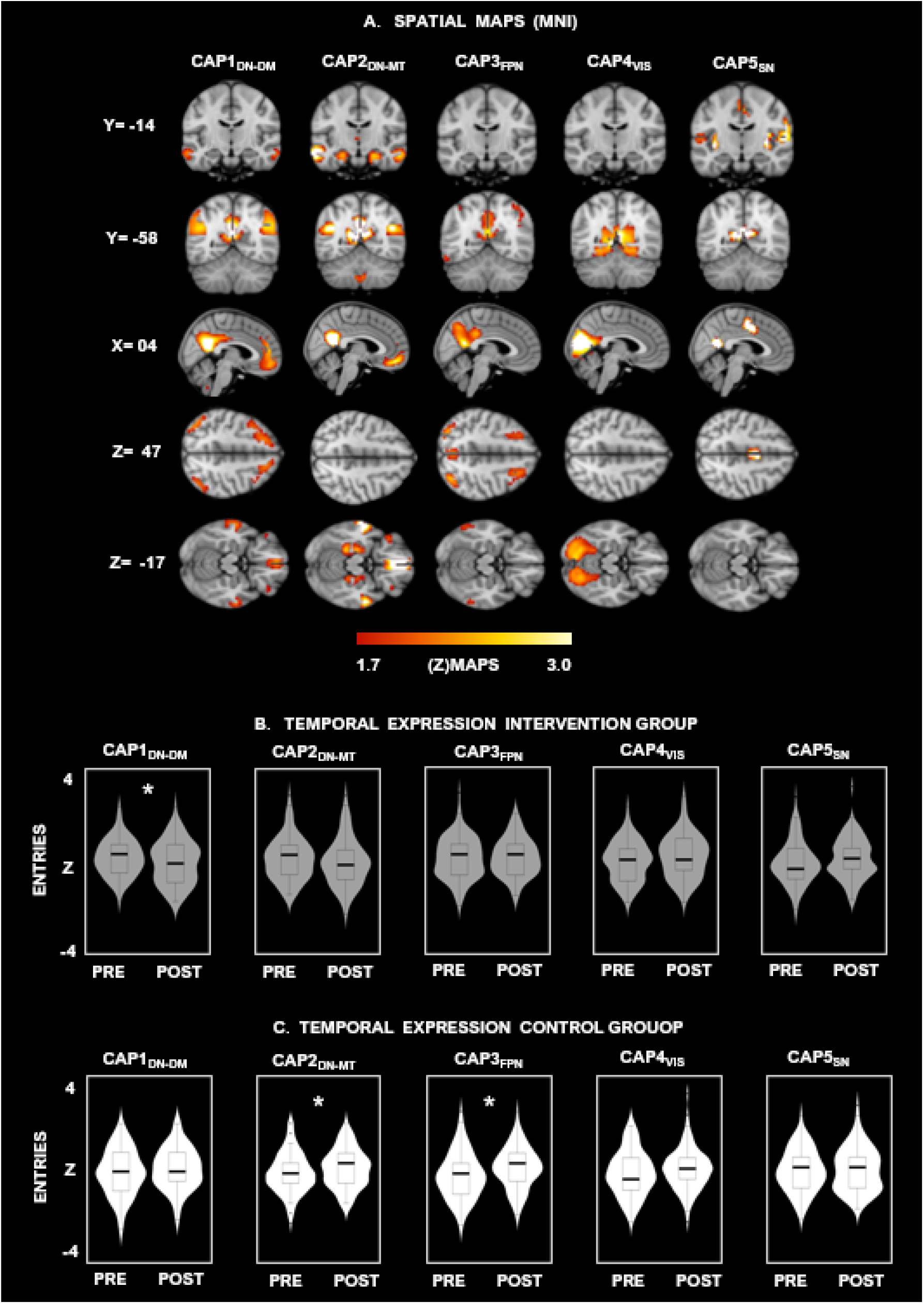
Brain maps depict the five co-activation patterns identified by the PCC-based co-activation patterns (CAPs) analysis. **A**. The spatial maps of each CAP are illustrated with a threshold of [z > 1.7 (p<.05); yellow color code]. Further anatomical information on peak regions is provided in **Tables S3, S4, S5, S6**, and **S7**, respectively. **B, C**. Violin plots depicting the median (horizontal black lines), the interquartile range (box markers), and the probability density (vertical histograms) of each CAP expression (entries) across groups [“INTERVENTION” (grey color), and “CONTROL” (white color)] and measurements collected before (“PRE”) and following (“POST”) the mindfulness-based intervention (MBI) or waiting period. Inferential statistics on the temporal entry rates are reported in **Table S1**. * p<0.05.

### Association of clinical measures with CAPs

Next, we assessed whether the entry rates of CAPs with greater response to MBI were associated with self-reported measures of stress, depression, and anxiety (i.e., STAI-S, STAI-T, BDI, and DASS scores) and whether these associations changed in response to i) the type of feedback in the MIST task [i.e., neutral, negative, and positive (**Figure 1B**), and ii) the mindfulness MBI intervention. We computed non-parametric Spearman coefficients (RHO). The probability of observing a non-zero RHO coefficient for each association (i.e., p values) was adjusted with a false discovery rate (FDR) for multiple testing (88), which is also optimal for controlling group dependency in nested data (89).

### Causal mediation of mindfulness trait on brain-behavior interactions

Lastly, we probed the impact of mindfulness training on the relationship between brain activity (i.e., CAPs) and clinical symptoms (i.e., STAI-S, STAI-T, BDI, and DASS). For this assessment, we quantified the causal mediation of the levels of acceptance and mindfulness (i.e., CAMM scores) on linear regressions between the brain CAPs and the clinical scores. The causal mediation approach (90) evaluates the causal role of a mediator, which may not be the case with conventional analysis. To give the average mediation effect a causal interpretation, it is critical to determine the impact of unmeasured pre-intervention or post-intervention confounders on the estimates of an average causal mediation e□ect [i.e., ACME. (91)]. The causal mediation analysis allows us to examine how sensitive the ACME estimates are to unmeasured confounders through the correlation between the residuals of the mediator (*e*_*m*_) and outcome regressions (*e*_*y*_):

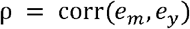

Where ρ represents the amount of pre-intervention mediator-outcome confounding necessary to result in no ACME. Namely, a larger p-value indicates robustness in the ACME estimate (92)].

## Results

### Dynamic functional connectivity (dFC)

We computed brain functional CAPs with the mid-frontal (MFG), inferior frontal gyrus (IFG), and posterior cingulate cortices (PCC) as seed regions of interest (**Figure 1E**) for pursuing our main goal: To unveil the effects of 8-week MBI on neural activity underlying the recovery from acute social stress in adolescents. Preliminary inspections showed satisfactory models’ compliance with the assumptions for linear regressions. The “consensus” algorithm indicated K=5 as the optimal number of solutions for the three seed regions (**Figure 1D**). Subsequently, the test-retest reliability of these networks (i.e., CAPs) was validated with CAPs from further k-mean solutions (k= 3, k= 4, k= 5, k= 6, k= 8, k= 12). The intercluster similarity analyses yielded high spatial consistency of the CAPs initially identified across different K-mean solutions. Namely, the five CAPs from the reference clustering (k=5) showed high overlap (mean=78%) with similar CAPs resulting from different k-mean solutions (**Figure 1C**). Results from the LMMs yielded nonsignificant differences between both groups (i.e., “intervention” and “control”) when the co-activation patterns (i.e., CAPs) derived from the MFG and IFG seed regions were compared before and following the mindfulness intervention (i.e., “pre”, and “post” measures). Likewise, no differences were detected when the type of feedback in the MIST was controlled (i.e., “neutral,” “negative,” and “positive”). Functional maps related to the bilateral MFG and IFG co-activation patterns (CAPs) are illustrated in **Figure S1** and **Figure S2**, respectively. Results regarding their temporal expression (i.e., entry rates) are reported in **Table S1**. Only the PCC co-activation patterns showed group differences.

### PCC-related dynamic functional co-activation patterns (CAPs)

Entry rates from three CAPs derived from the posterior cingulate cortex (i.e., PCC) differed between control and intervention groups before and following the mindfulness intervention (see “pre” and “post” MBI measurements in **Figure 2)**. However, such differences were not statistically significant when the type of feedback in the MIST was controlled (i.e., neutral, negative, and positive. See results in **Table S1**). Assessments of the model’s integrity yielded adequate goodness of fit in the PCC-CAPs. However, the model related to PCC-CAP1 reported better performance compared to further CAPs. Results from model comparisons are reported in **Table S2** and **Figure S3A**. Also, posterior predictive checks yielded consistency between the maximum likelihood parameter estimation (MLE) and the real data (**Figure S3B**). Furthermore, the inspection of influential observations (i.e., outliers. **Figure S3C**), normality of residuals (i.e., normal distribution. **Figure S3D**), collinearity among predictors (**Figure S3D**), and the homogeneity of variance (i.e., homoscedasticity. **Figure S3F**) confirmed an optimal estimation of the modeled parameters.

This first CAP overlapped the dorsal-medial subsystem of the default network [CAP1_DN-DM_ (93)] **Figure 2A**. See peak coordinates in **Table S3**], including the bilateral superior and mid-frontal gyri, medial prefrontal cortex (MPFC), posterior cingulate cortex (PCC), precuneus, and the bilateral mid-temporal gyri. Entries of this CAP1_DN-DM_ (i.e., the number of times that the brain entered this state of co-activation) differed significantly between groups (i.e., “intervention” vs. “control”) and visits [i.e., “pre” mindfulness intervention (MBI) vs. “post” MBI (F(1, 318.09)=4.87. P= 0.03. *η*2= 0.02)]. Such difference was triggered by a decrease of CAP1_DN-DM_ entries in the intervention group “post-MBI” compared to _CAP1DN-DM_ entries in “pre-MBI” [βintervention “pre”= 0.17; βintervention “post”= -0.14. P_FDR_(CI 0.01; 0.56)= 0.06. D_COHEN_=0.36)]. Conversely, the control group reported an increase in the CAP1_DN-DM_ entries following the MBI compared to the baseline (“pre”) measurements. Notably, this increase was a trend [βcontrol “pre”= -0.05; βcontrol “post”= -0.03. P_FDR_(CI -0.47; 0.08)=0.17. D_COHEN_= -0.20. **Figure 2B**, full statistics in **Table S1**)].

A second CAP encompassed regions from the medial temporal subsystem of the DN (i.e., CAP2_DN-MT_), including the hippocampus, the parahippocampal gyri, posterior cingulate areas, fusiform gyri, mid-temporal gyri, and the anterior cingulate cortex. (**Figure 2A;** see peak coordinates in **Table S4**). Entry rates of the control group featured by this CAP2_DN-MT_ increased in the “post” measurement compared to the baseline measures. [βcontrol “pre”= -0.15; βcontrol “post”= -0.08. P_FDR_(CI -0.54; 0.01)=0.05. D_COHEN_= -0.27. **Figure 2B**. Full statistics in **Table S1**)]. The third CAP overlapped areas comprising the Frontoparietal Network, including the superior parietal lobes, angular gyri, cingulate gyri, and mid and superior frontal gyri (i.e., CAP3_FPN_). CAP3_FPN_ entries differed between groups (i.e., “intervention” vs. “control”) and measurements (i.e., “pre” and “post” mindfulness training) [*F*(1, 318.25)= 3.23. *P=* 0.07. *η*^*2*^= 1.00]. Post-hoc analyses revealed an increase of CAP3_FPN_ entries in the “post” measurement of the control group [βcontrol “pre”= -0.30; βcontrol “post”= -0.07. P_FDR_(CI -0.69; 0.14)=0.00. D_COHEN_= -0.41]. **Figure 2B**, full statistics in **Tables S1**)]. A fourth CAP implicated regions of the visual system [CAP4_VIS_(94)], comprising lingual and fusiform gyri, cuneus, calcarine, occipital mid, occipital superior, and posterior parietal areas (**Figure 2A**; see peak coordinates in **Table S6**). Finally, a fifth co-activation pattern encompassed regions associated with the salience network (CAP5_SN_). Core regions of this CAP included bilateral anterior insula and anterior midcingulate cortices, the thalamus, supramarginal and precuneal areas (**Figure 2A**; see peak coordinates in **Table S7**). The entry rates of these CAPs showed no significant differences in terms of groups and visits [CAP4_VIS_: (*F*(1, 317.70)= 0.10 / P= 0.75. *η*^*2*^= 0.01). CAP5_SN_: (*F*(1, 317.76)= 0.55. P= 0.46. *η*^*2*^= 0.02). **Figure 2B**, full statistics in **Tables S1**)].

Overall, the multilevel linear mixed modeling (LMM) revealed brain systems with different responses to our mindfulness-based intervention (MBI), regardless of the type of stress elicited in the MIST paradigm (i.e., neutral, negative, or positive feedback. See **Figure 1B**). PCC-CAP1_DN-DM_ recorded from the “intervention” group decreased following the mindfulness-based intervention (i.e., MBI). Conversely, PCC-CAP2_DN-TM_ and PCC-CAP3_DN-DM_ presented greater entry rates after the waiting period in the control group. Next, we explore the interaction between these brain CAPs and clinical/behavioral indicators of anxiety and depression.

### Relationship of brain CAPs with levels of anxiety and stress

Next, we assessed (i) whether these brain CAPs with differential responses to MBI (i.e., CAP1_DN-DM_, CAP2_DN-MT_, and CAP3_FPN_) were associated with clinical scores related to anxiety and depression and (ii) whether those brain-behavior correlations change as a function of the type of stress elicited in the MIST feedback. In accordance with the previous multilevel LMMs, assessments of non-parametric correlations yielded nonsignificant associations between these variables in either group when the type of MIST feedback was controlled. The results are reported in **Figure S4**. Conversely, the number of entries into the brain CAP1_DN-DM_ significantly correlated with the clinical scores when the type of MIST feedback was not controlled (**Figure 3A**). Associations with the strongest magnitude were observed preceding the mindfulness training [Interventions “pre” measure: CAP1_DN-DM_ - anxiety-state (STAI) rho(93)= 0.26; CI 95%(0.05; 0.44); Interventions “pre” measure: P_FDR_= 0.02. CAP1_DN-DM_ - anxiety-trait (STAI-T) rho (93)= 0.31; CI 95%(0.11; 0.49); P_FDR_= 0.00. Interventions “pre” measure: CAP1_DN-DM_ - depression (BDI) rho (93)= 0.28; CI 95%(0.07; -0.46) ; Controls “pre” measure: P_FDR_= 0.01. CAP1_DN-DM_-depression (DASS-D) rho(105)= -0.25; CI 95%(−0.43; -0.06); P_FDR_= 0.02. **Figure 3A**]. Next, we tested the impact of mindfulness on the associations between CAP1_DN-DM_ and behavioral scores related to psychological distress in both groups.

**Figure 3.**
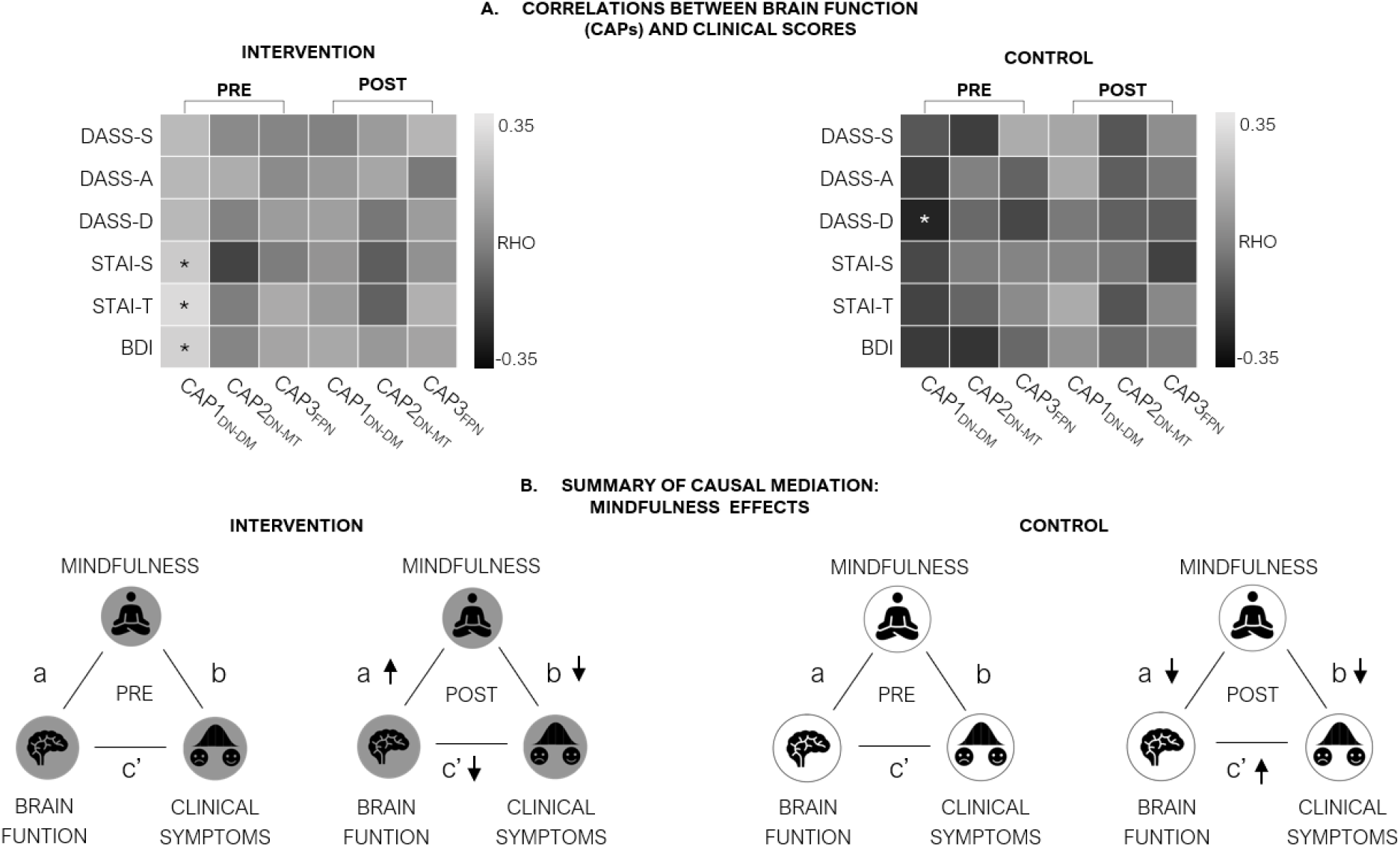
**A**. Non-parametric Spearman associations (RHO) between brain activity and pre-post questionnaire items and state anxiety score. The clinical scores portraying the stress and anxiety status of our participants were screened twice before (i.e., “pre” measurement) and following the mindfulness intervention (i.e., “post” measurement), respectively. **B**. Results summary of mediation analyses. The path [c’ (↓), “POST” diagram] represents a decreased magnitude in the link between brain function (i.e., CAP1DN-DM) and self-reports of depression and anxiety (i.e., STAI-S, STA-T, BDI, DASS-D) in response to mindfulness training in the intervention group. The path [a (↑). “POST” diagram] represents a change in the direction, from (−) to (+), in the link between brain function (i.e., CAP1DN-DM) and mindfulness trait (i.e., CAMM). The path [b (↓), “POST” diagram] represents a strengthening in the negative correlation between mindfulness trait and clinical symptoms. The direction of the same associations was the opposite in controls. E.g., a stronger positive correlation between brain function and clinical symptoms in the “POST” diagram. Values of the causal mediation paths are described in Tables S5 and S6.

### Causal mediation of mindfulness trait on brain-behavior interactions

We then probed the causal mediation of mindfulness on the relationship between brain activity and psychological distress using a direct measure of trait mindfulness (CAMM). Consistent with the nonparametric correlations, preliminary linear regressions (i.e., total effects in mediation models). revealed associations between brain CAP1_DN-DM_ and self-reported scores related to depression (i.e., BDI, DASS-D) and anxiety (i.e., state and trait STAI). Notably, these associations changed in both groups when “pre” and “post” measurements were compared. In the intervention group, associations between CAP1_DN-DM_ and the clinical symptoms decreased as a function of MBI (see total effects in **Tables 2, 3**, and summary in **Figure 3B** left). This effect was causally mediated by CAMM scores. More specifically, the “a” estimate (i.e., β of CAP1_DN-DM_ ∼ CAMM) shifted in a positive direction in response to the MBI mindfulness intervention (see a(β) pre-measurement and a(β) post-measurement in the path diagrams of **Figures S5, S6**, and summary of effects **in Figure 3B** left). In contrast, “b” estimates (i.e., β of CAMM ∼ clinical scores) reported a negative shift in response to MBI (see b(β) pre-measurement and b(β) post-measurement in diagrams of **Figures S5, S6**, and summary of effects in **Figure 3B** left). The inverse pattern was observed in the control group. Namely, the total effects of CAP1_DN-DM_ entries on clinical symptoms shifted from negative estimates in the “pre” measurement to positive estimates in the “post” measurement. See total effects in **Tables 2**,**3** and a summary of effects in **Figure 3B** right). Likewise, β estimates of the regression CAP1_DN-DM_ ∼ CAMM (i.e., “a” path) flipped negatively in the “post” measurement (see **Figures S5, S6**, and summary of effects in **Figure 3B** right). In addition, post-hoc sensitivity analyses revealed no evidence of unmeasured confounders influencing the average causal mediation (i.e., the ACME estimates) implemented in both groups. Reports from the sensitivity assessments are found in Figure **S7**.

**Table 2.**
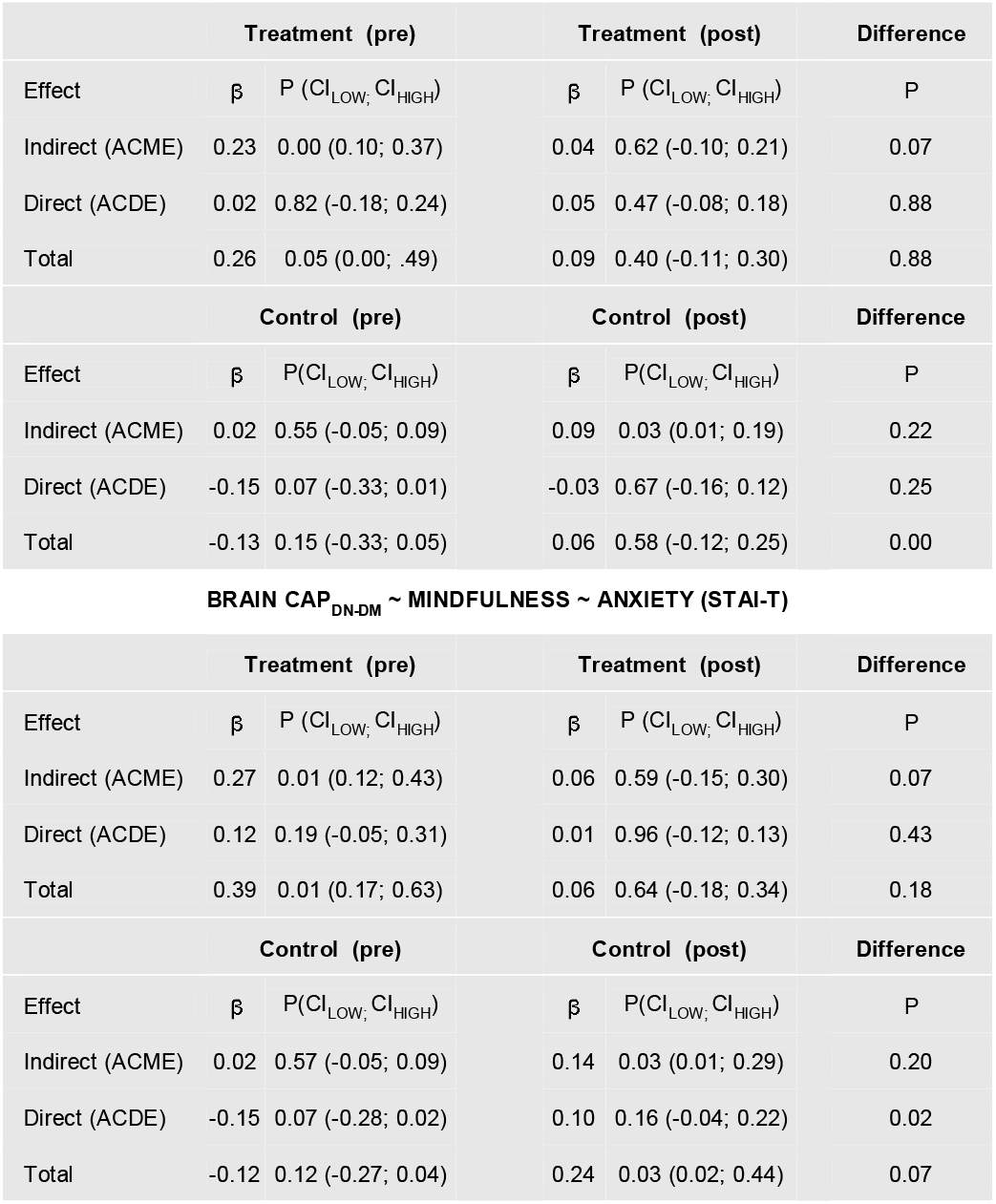
Results of multilevel causal mediation analyses. Average causal mediation (ACME), average causal direct (ACDE) and total effects of mindfulness (i.e., CAMM) on the relationship between brain function (i.e., CAP1_DN-DM_) and anxiety states (i.e., STAI-S and STAI-T). The difference between the effects before mindfulness (i.e., “pre”) and following mindfulness (training (i.e., “post”) was quantified through the probability (P) under the assumption of no difference.

**Table 3.**
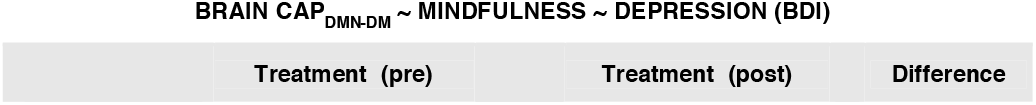
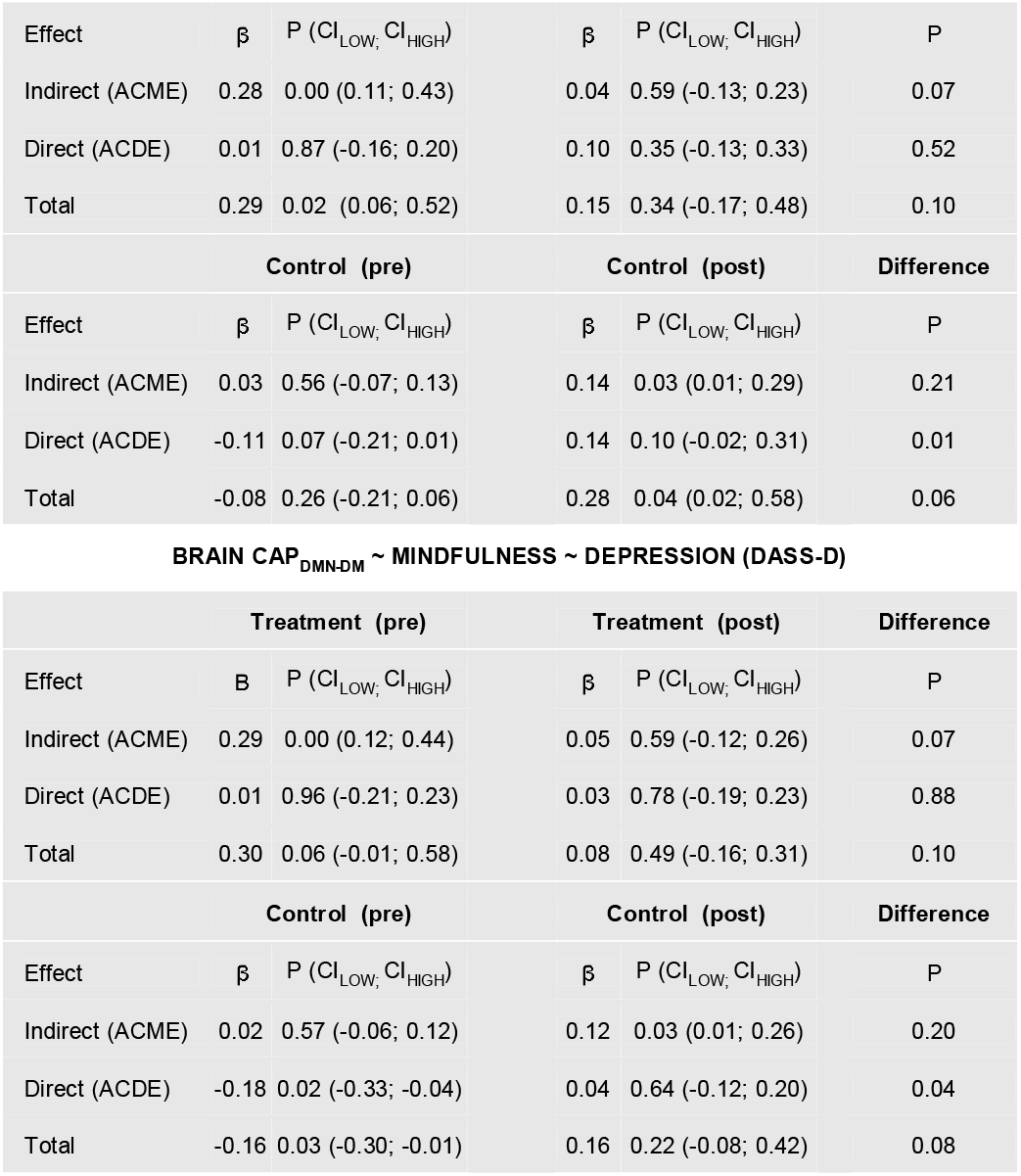
Results of multilevel causal mediation analyses. Average causal mediation (ACME), average causal direct (ACDE) and total effects of mindfulness (i.e., CAMM) on the relationship between brain function (i.e., CAP1_DN-DM_) and depressive symptoms (i.e., BDI and DASS-D). The difference between the effects before mindfulness (i.e., “pre”) and following mindfulness (training (i.e., “post”) was quantified through the probability (P) under the assumption of no difference.

## Discussion

We evaluated the impact of mindfulness on the neural response to psychosocial stressors in non-clinical adolescents. Specifically, we explored whether (i) the dynamics of functional brain activity exhibits specific patterns (i.e., CAPs entries) in response to mindfulness intervention. And (ii), whether the temporal expression of these CAPs unveils specific brain responses to different types of psychosocial stress (i.e., neutral, negative, and positive). The obtained results only confirmed our first hypothesis, and they will be discussed next.

### Brain CAPs linked to mindfulness trait

In line with our first hypothesis, the results yielded changes in the brain dFC as a function of mindfulness training. However, we did not confirm our second hypothesis. Namely, dFC changes related to the mindfulness intervention were not sensible to the specificity of the psychosocial stressors with neutral, negative, or positive affective valence. Brain co-activation patterns (i.e., CAPs) with greater response to MBI encompassed dorsal-medial areas of the DN (i.e., CAP1_DN-DM_). In the intervention group, the number of times the brain “entered” into the CAP1_DN-DM_ state (i.e., entry rates) was higher before (i.e., “pre”) MBI in comparison to the entry rates “post” MBI. Furthermore, the same CAP1_DN-DM_ entry rates were significantly linked to fewer indices of mindfulness trait (i.e., CAMM), also in “pre” MBI. This anticorrelation between brain dynamics and mindfulness trait drastically shifted after mindfulness training (**Figure 3B**). In line with previous research, our results suggest that MBI hinders the effect of acute stress in adolescents at the brain level. This interpretation is also grounded in the opposite pattern observed in controls (**Figure 3B**). Reduced CAP1_DN-DM_ dynamics portraying the effects of MBI on stress response is also aligned with previous reports which, in contrast to our results, reported structural and functional hyperconnectivity between DN areas in adolescents experiencing acute stress (28), posttraumatic stress symptoms (95), and depression (96,97).

### Relationship of brain CAPs with clinical symptoms

One further result described changes in the relationship between CAP1_DN-DM_ entries and anxiety and depression symptoms. Greater entry rates of CAP1_DN-DM_ were significantly correlated to higher anxiety and depression rates before mindfulness training (i.e., “pre” MBI). The direction and strength of these associations also switched in response to MBI only in the intervention group (**Figure 3B**). Extensive research links enhanced activity in DN underlying the experience of acute stress, anxiety, and depression (98–101). However, this relationship might not be causal due to potential influences from uncontrolled factors that might play important roles (e.g. levels of neurotic trait, reaction to MRI scanning, etc.). Interestingly, the link between CAP1_DN-DM_ and clinical outcomes weakened after MBI. Therefore, we assessed whether the mindfulness trait plays a deterministic role in this brain-behavior relationship.

### Mindfulness trait causally mediates the impact of acute stress in the functional brain

We examined how mindfulness training (MBI) affects the way the brain responds to social and psychological stress in healthy teenagers. Our study found that MBI reduced the number of times the brain entered a state characterized by higher connectivity between dorsal medial DN areas. The reduced number of entries to this CAP1_DN-DM_ state was associated with lower levels of anxiety and depression symptoms. In contrast, neither this brain pattern nor the association with symptoms was observed in the control group. Additionally, results from this study support a neurocognitive model in which mindfulness trait causally mediates the interaction between brain dynamics and clinical symptoms. Reduced entry rates into brain CAP1_DN-DM_ state are linked to increased mindfulness trait and reduced scores of depression and anxiety symptoms. Therefore, the number of entries into the CAP1DN-DM brain state predicts the presence of clinical symptoms, and this association is causally mediated by the mindfulness trait. This causal mediation model aligns and brings together previous findings in which individuals exposed to acute stress endorsed greater mind wandering (102), which in turn has been linked to DN hyperconnectivity (103). Furthermore, a recent pilot study showed reduction in DN activation in response to mindfulness-based fMRI neurofeedback in adolescents with major depression (104).

## Conclusions

We analyzed fMRI data from an RCT to explore the impact of MBI on dFC responses related to psychosocial stress recovery in adolescents. MBI reduced the number of entry rates of CAP1_DN-DM_ state and its associations with symptoms of anxiety and depression. Here, we proposed a brain-behavior model describing how mindfulness trait decreases the activity of specific DN areas and reduces psychological distress in adolescents. The present study postulates a novel dFC metric of mindfulness trait by quantifying the entries into a particular brain state (e.g., CAP1_DN-DM_). Globally, our findings add new evidence regarding the neural basis of mindfulness trait and their potential as biomarkers of stress resilience and vulnerability.

## Limitations and future directions

Our CAPs analyses could not differentiate the neural response to the affective valence of our psychosocial stressors. The implementation of alternative dFC markers might shed new insights into how the brain processes different types of stress-laden stimuli. In addition, future research might also assess multimodal data including fMRI and neuroendocrine recordings to better understand the biology of stress reactivity and its modulation through MBI. Lastly, implementing the same study with long-term follow-up in adults and the elderly will advance our understanding of stress vulnerability, reactivity, and resilience across the human life span.

## Data availability

Raw data supporting the findings of this study will be available following the brain imaging data structure (i.e., BIDS(105)) at https://layerlab.unige.ch/#/home, in full accordance to the FAIR principles for scientific data management(106).

