## Supplementary material for "Mindfulness training impacts brain network dynamics linked to stress response in young adolescents": Suppl. material

| **Authors** | Julian Gaviria *^a, b, c, d^. ORCID ID: 0000-0002-4266-1371  Zeynep Celen ^d^. ORCID ID: 0000-0000-0002-1227-6587  Mariana Magnus Smith ORCID ID: 0000-0000-0000-0000  Lucas Peek ^e^. ORCID ID: 0000-0000-0000-0000  Soraya Brosset ^e^. ORCID ID: 0000-0000-0000-0000  Patrik Vuilleumier ^e^. ORCID ID: 0000-0002-8198-9214  Dimitri Van De Ville ^f, g^. ORCID ID: 0000-0002-2879-3861  Arnaud Merglen ORCID ID: 0000-0000-0000-0000  Paul Klauser ORCID ID: 0000-0000-0000-0000  Camille Piguet ^d^. ORCID ID: 0000-0002-8198-9214  ^a^ Department of Psychiatry, ^b^ Amsterdam Neuroscience, Amsterdam UMC; ^c^ Vrije Universiteit, Amsterdam. The Netherlands. ^d^ Department of Psychiatry; ^e^ Laboratory for Behavioral Neurology and Imaging of Cognition, department of Basic Neurosciences; ^f^ department of radiology and medical informatics, University of Geneva. Switzerland; ^g^Medical Image Processing Lab, Neuro-X Institute, Ecole polytechnique fédérale de Lausanne (EPFL). Switzerland**;** |
| --- | --- |
| ***Correspondence** | Julian Gaviria.; |
| **Keywords** | Brain networks; mindfulness; dynamic functional connectivity (dFC); depression; stress; adolescents; anxiety, stress |

### **Supplementary methods**

#### **FMRI data acquisition**

Neuroimaging data were collected using a 3T Magnetom TIM Trio scanner (Siemens, Germany) at the Brain and Behavior Laboratory at the Faculty of Medicine of the University of Geneva (Switzerland). We used a 32-channel head-coil. Dynamic functional connectivity of the resting state was computed from Blood Oxygenation Level Dependent (BOLD) contrast from a T2*-weighted echo-planar sequence (EPI). 2160 functional volumes (1080 per visit) of 60 axial slices each (TR/TE/flip angle = 2100ms/25ms/60°, FOV=212 mm, resolution=106×106, isotropic voxels of 2 mm^3^, distance factor 10%) were acquired. Furthermore, we collected a high-resolution T1-weighted anatomical image [TR/TI/TE/flip angle=1900ms/900ms/2.27ms/9°, FOV=256mm, resolution=256×256, slice thickness=1mm^3^, 192 sagittal slices, phase encoding direction= Anterior-Posterior (AP). Our multiband sequence included 4 dummy scans (~ 5s) at the beginning of the fMRI scanning.

#### **Preprocessing for co-activation pattern (CAP) analysis**

Following Bolton et al.,^14^ standard image preprocessing procedures were applied using the DPABI toolbox^15^. Time-series from each voxel in the white matter and cerebrospinal fluid, plus the six affine motion parameters from realignment (also including their respective first-order derivatives), were used as nuisance variables to be regressed out from the data. No global signal regression (GSR) was carried out given the absence of agreement in the field in this regard^16^. Although some studies support this approach^17,18^, others suggest that GSR does not affect the CAP extraction procedure^19^ nor measures of anticorrelation among intrinsic functional brain networks^20^. Furthermore, the data were band-pass filtered between 0.01 and 0.10 Hz. All image volumes with frame-wise displacement above 0.5 mm were discarded, as well as subjects with more than 30% of scrubbed frames^21^.

#### **FMRI Task**

The task was presented in two sessions, with 6 runs in each session. The order of the runs was pseudorandomized into 6 lists. The experimental runs consisted of an information screen (5sec) signalling the session will be evaluated; followed by a series of 5 time-limited challenging mental calculations, followed by a positive or negative feedback with a social ranking (high for positive, low for negative) last lasted for 8 sec. After the feedback, participants had a 90 sec recovery period (i.e., resting state), where an information screen for 5 sec was presented which instructed the participants to close eyes and rest, then the screen turned black for 85 sec. Participants were signaled the end of the rest period by 2 flashes of light and a sound. There were 8 experimental runs, and 4 control runs where the information screen signaled the session will not be evaluated, the calculations were simple and the feedback screen had a neutral feedback with no ranking. The recovery period information screen and time was the same for all runs. The whole task lasted around 25 minutes. The participants’ eye movements were observed during the experiment to make sure they close their eyes during the recovery period.

#### **Linear mixed modelling**

We performed a 2 × 2 x 3 LMM-based factorial analyses denoted by the following R-based formula syntax:

entries ∼ group * measurement * condition +(1 |subject ),

where the ﬁxed factor “group” comprised two levels corresponding to assigned cohorts in the randomization (i.e., “treatment” and “control”). The second fixed factor “measurement” included the measurements recorded in relation to the MBI mindfulness intervention (i.e., “pre” and “post”). The fixed factor “condition” comprised the type of feedback in the MIST task preceding the resting periods (i.e., “neutral”, “negative”, and “positive”). Individual participants (i.e. “subject”) were modeled as a random factor, and the “entries” were the dependent variable, computed for each CAP. Preliminary inspections of the models examined the compliance with the assumptions for linear regressions in terms of influential observations (i.e., outliers), homogeneity of variance, normality of residual, and collinearity. Additionally, posterior predictive checks assessed the consistency between the maximum likelihood parameter estimation (MLE) and the empirical data.

### **Supplementary Figures**

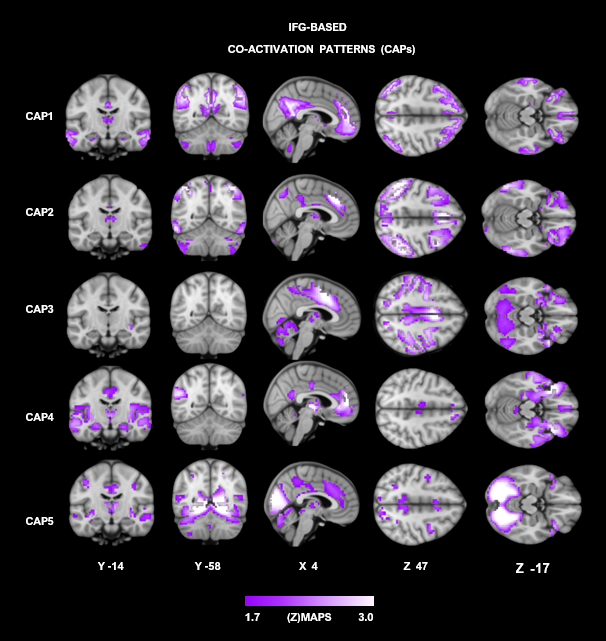

**FigureS1**. Brain maps depict the five co-activation patterns derived from the bilateral med frontal gyri (MFG). **A.** The spatial maps of each CAP are illustrated with a threshold of [z > 1.7 (p<.05); purple color code]. Results of the inferential statistics on the entry rates of these MFG-based CAPs are described in **Table S1**.

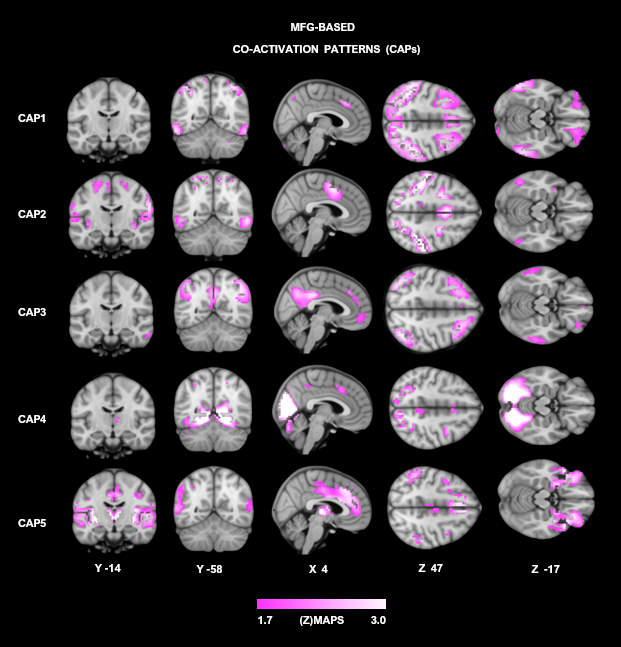

**FigureS2**. Brain maps depict the five co-activation patterns derived from the bilateral med frontal gyri (MFG). **A.** The spatial maps of each CAP are illustrated with a threshold of [z > 1.7 (p<.05); purple color code]. Results of the inferential statistics on the entry rates of these MFG-based CAPs are described in **Table S1**.

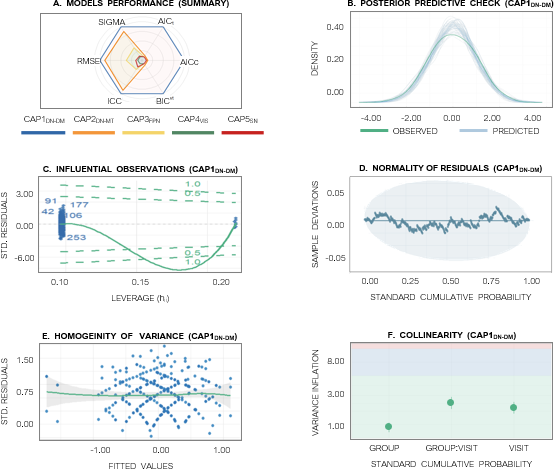

**Figure S3.** Model integrity assessments. The multilevel linear mixed models (LMM) implemented to examine the temporal fluctuation of co-activation patterns (CAPs) derived from the posterior cingulate (i.e., PCC) region of interest (i.e., ROI) yielded adequate of goodness-of-fit across groups (i.e., “treatment” and “control”) and measurements (i.e., “pre” and “post” mindfulness-based intervention). **A.** CAP1_DN-DM_ reported the best performance compared to further CAPs. A quantitative ranking of our model performance (%) is described in **Table S2**. **B.** Posterior predictive checks yielded consistency between the maximum likelihood parameter estimation (MLE) and the empirical data from CAP1_DN-DM_ (i.e., the entry rates). Furthermore, the inspection of influential observations [i.e., outliers (**C**)], normality of residuals [i.e, normal distribution (**D**)], collinearity among predictors (**E**), and the homogeneity of variance [i.e., homoscedasticity (**F**)] confirmed an optimal estimation of the modelled parameters for the models. As a representative sample, Evaluation of CAP1_DN-DM_ is represented here.

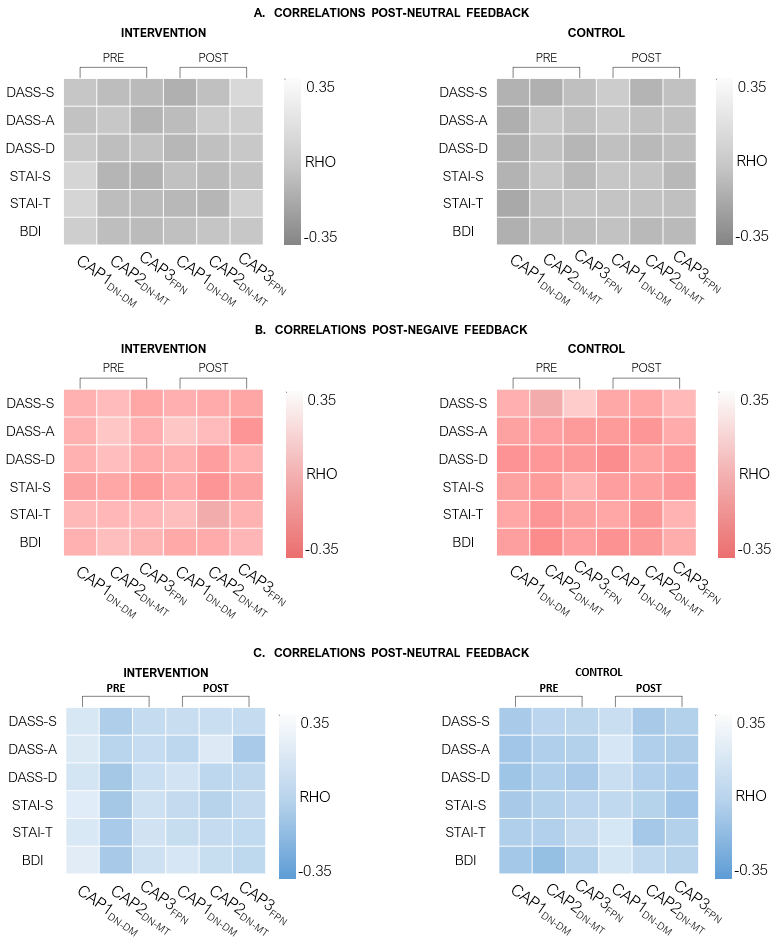

**Figure S4.** Non-parametric Spearman associations (RHO) between brain activity and clinical scores. The clinical scores portraying depression and anxiety status of our participants included the State-Trait Anxiety Inventory for Children (STAI-T, STAI-S); the level of depressive symptoms was assessed by the Beck Depression Inventory (BDI) and the Depression and Anxiety Stress Scale (DASS). Y-axis in matrices. These self-reports were screened before (i.e., “pre” measurement) and following the mindfulness intervention (i.e., “post” measurement) respectively. The brain activity was represented by the entry rates of CAPs with greater response to the mindfulness-based intervention: CAP1_DN-DM_, CAP2_DN-MT_, and CAP3_FPN_ (x-axis in matrices). The brain CAPs were recorded following the MIST stimuli with neutral (A), negative (B), and positive feedback (C) respectively. None of the correlations reported significantly strong magnitude.

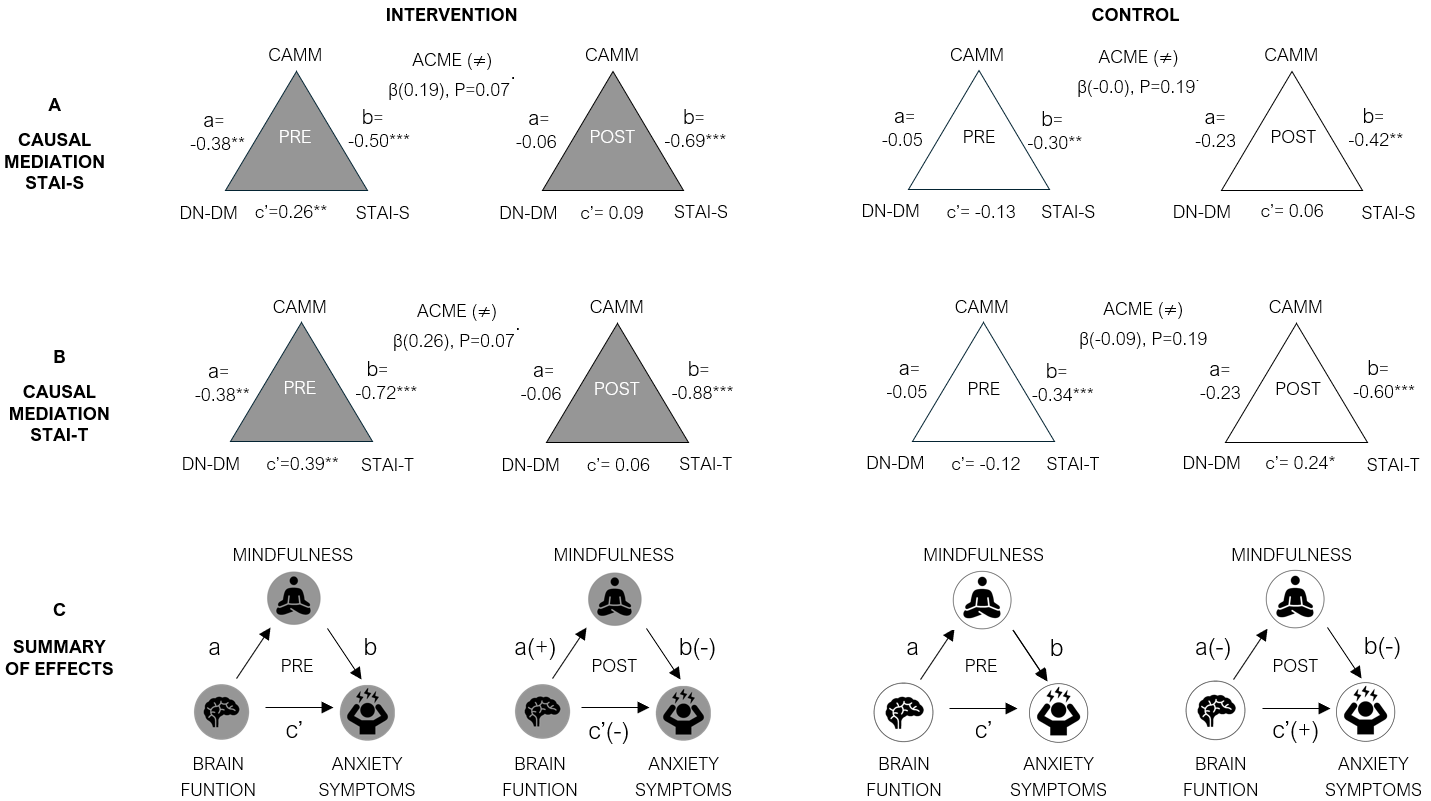

**Figure S5.** Causal mediation analysis. Path diagram for the causal mediation of mindfulness (i.e., CAMM scores) on the relationship between brain function (i.e., CAP1_DN-DM_) and symptoms related to anxiety [i.e., STAI-S(**A**), STAI-T(**B**). **^.^**p < 0.10. *p < 0.05. **p < 0.01. ***p < 0.001. The average causal mediation effect (i.e., ACME) was stronger in treatment group before the MBI mindfulness intervention. Conversely, the same ACME difference was significant in controls when following MBI. **C**. Summary. The results indicate a decreased magnitude in the link between brain function (i.e., CAP1_DN-DM_) and anxiety scores (i.e., STAI-S, STAI-T) in response to mindfulness training [see c’ (–) in the treatment group, “POST” diagram). The causal mediating role of mindfulness is featured by positive associations between brain function and indices of dispositional mindfulness skills [i.e., CAMM. See path “a(+)” in “POST” diagram].

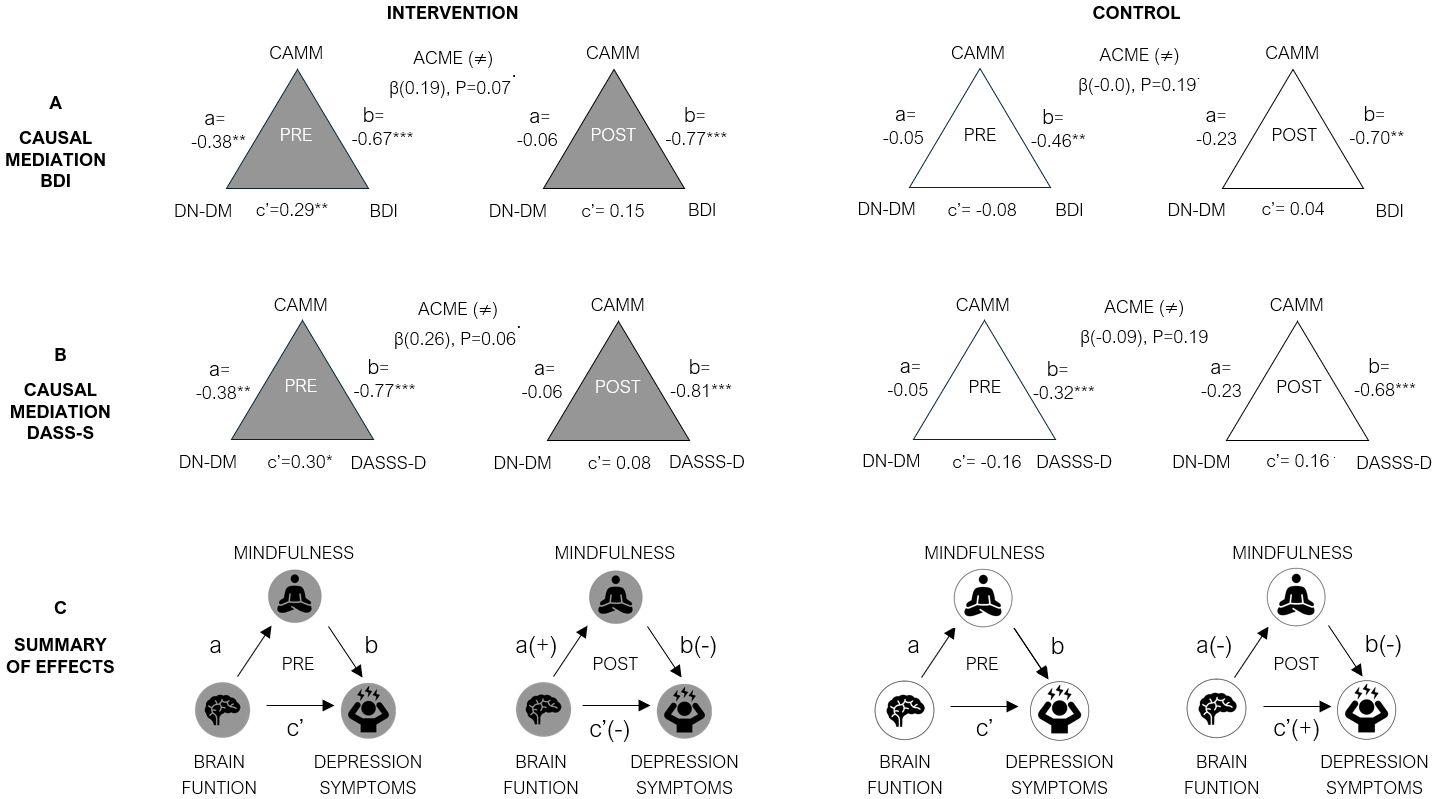

**Figure S6.** Causal mediation analysis. Path diagram for the causal mediation of mindfulness (i.e., CAMM scores) on the relationship between brain function (i.e., CAP1_DN-DM_) and symptoms related to depression [i.e., BDI (**A**), DASS-D (**B**). **^.^**p < 0.10. *p < 0.05. **p < 0.01. ***p < 0.001. The average causal mediation effect (i.e., ACME) was stronger in treatment group before the MBI mindfulness intervention. Conversely, the same ACME difference was significant in controls when following MBI. **C**. Summary. The results indicate a decreased magnitude in the link between brain function (i.e., CAP1_DN-DM_) and depression scores (i.e., BDI, DASS-D) in response to mindfulness training [see c’ (–) in the treatment group, “POST” diagram). The causal mediating role of mindfulness is featured by positive associations between brain function and indices of dispositional mindfulness skills [i.e., CAMM. See path “a(+)” in “POST” diagram].

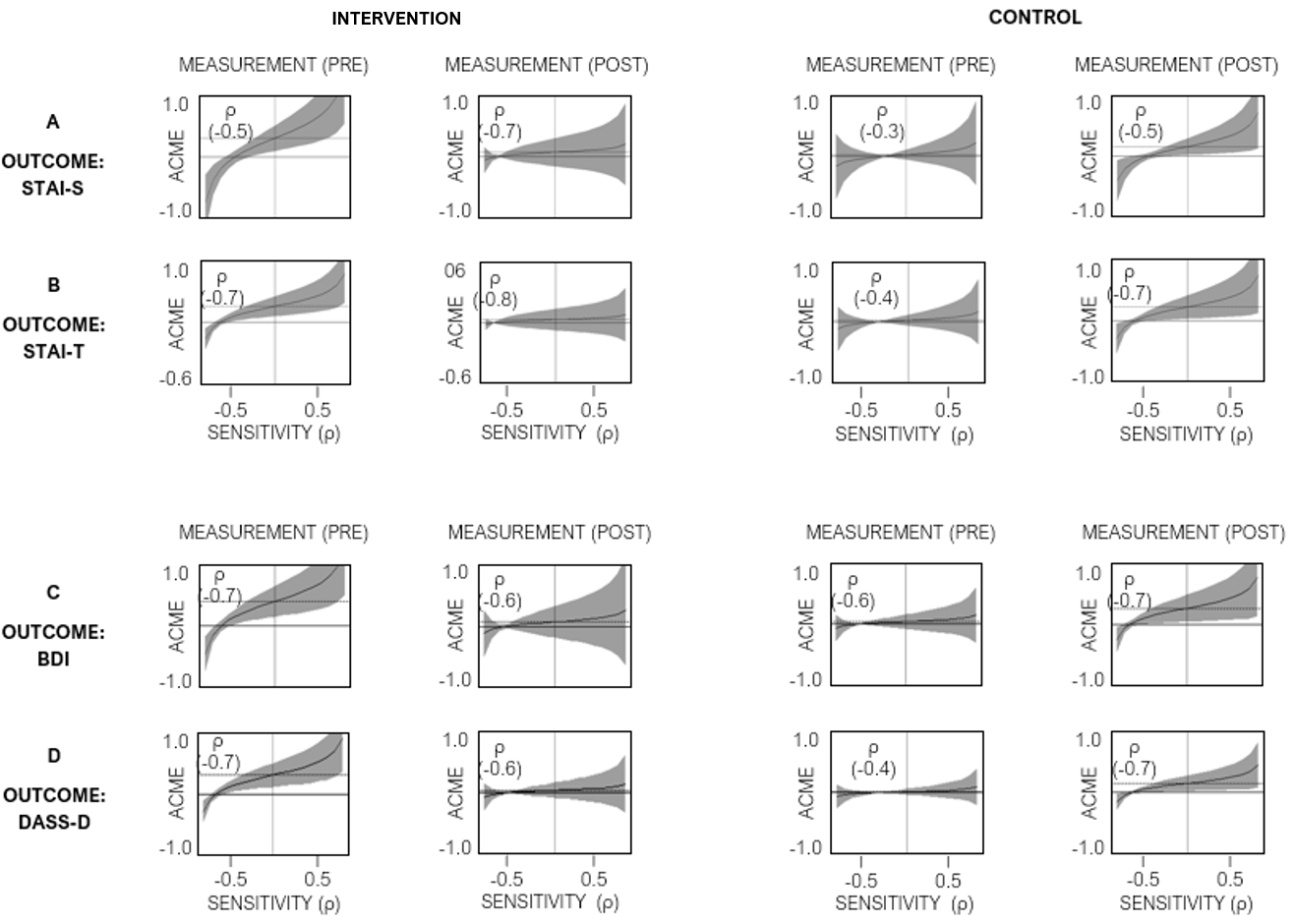

**Figure S7.** Sensitivity analyses. Sensitivity analyses implemented in the causal mediation models including self-reports of anxiety (**A**,**B**) and depression (**C**,**D**) revealed no evidence of unmeasured confounder. The dashed lines represent the estimated ACME. The gray areas represent the 95% confidence interval for the mediation effects at each value of the sensitivity parameter (ρ). Correlations between the residuals of the mediator and outcome regressions with coefficients below ρ =-0.24 indicate robust ACME values in relation to unobserved confounders (Keele et al., 2015, Imai et al 2010).

### **Supplementary Tables**

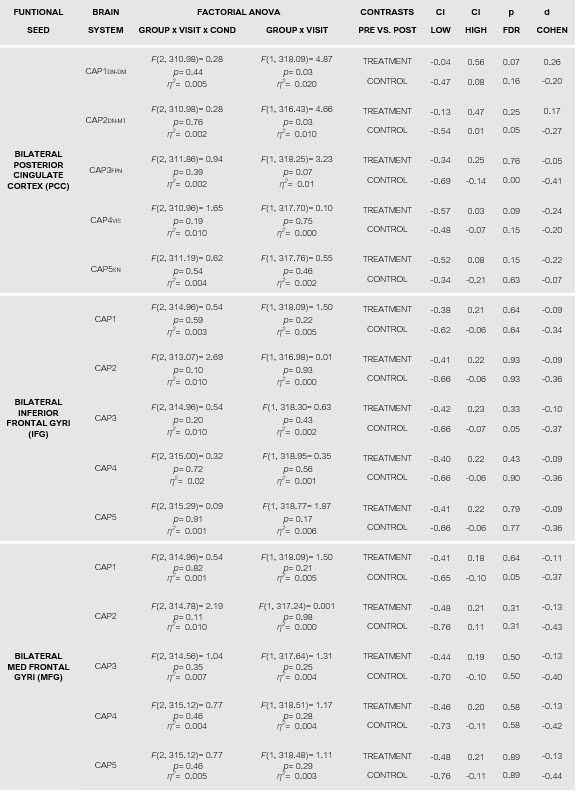

**Table S1.** Temporal expression of seed-based functional co-activation patterns (CAPs). The temporal entries of CAPs derived from posterior cingulate (PCC), medial frontal (MFG), and inferior frontal areas (IFG) were assessed through multilevel factorial models. The participants were modelled as random factor (i.e., repeated measures) and the factors “GROUP” (“treatment” and “control”), “VISIT” (“treatment” and “control”), and “CONDITION” (“neutral”, “negative”, and “positive”) as fixed variables. CAPs linked to the MFG and IFG brain seeds did not respond to our experimental manipulation. Conversely, CAP1, CAP2, and CAP3 derived from the PCC seed region showed different entry rates across groups (i.e., “treatment” and “control”) and measurements before (i.e., “pre”) and following (i.e., “post”) the mindfulness intervention. The percent of partial variance explained η^2^expresses the effect size of the main effects and interactions. Post-hoc pairwise least squared means contrasts (EMMEANS) driving the main effects and interactions are listed in the right-hand columns. Significance indices (*p_FDR_*) are adjusted for multiple testing under dependency. Effects size (*d*_Cohen_) corresponds to the effect size of the assessed contrasts.

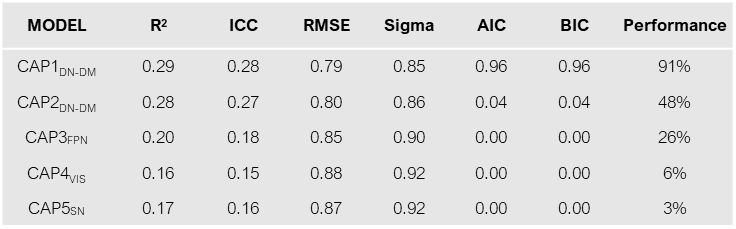

**Table S2.** General index of model performance (%). Performance of the linear mixed models were assessed through different metrics including the coefficient of determination (R^2^), intraclass correlation coefficient (ICC), root-mean-square error (RMSE), the residual standard error (sigma values), Akaike’s Information Criteria (AIC), and the Bayesian information criteria (BIC). These metrics comprise our index of model performance (%). Our Geneal index of model performance revealed a large difference between CAPs. The model related to CAP1_DN-DM_ reported the best performance. See also graphical summary in the first panel of **Figure S3**.

**CAP1_DN-DM_**

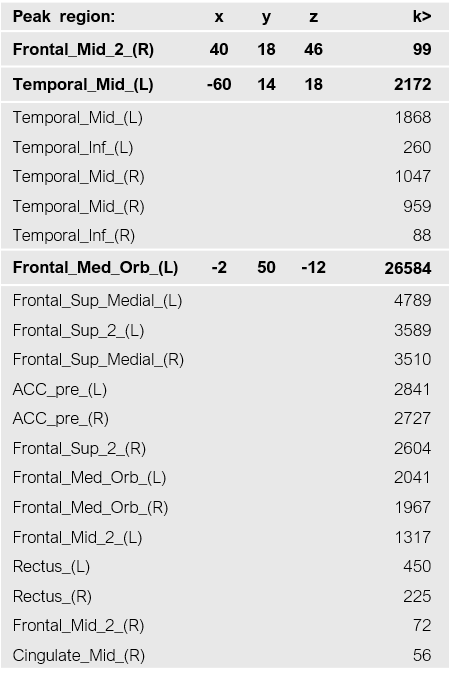

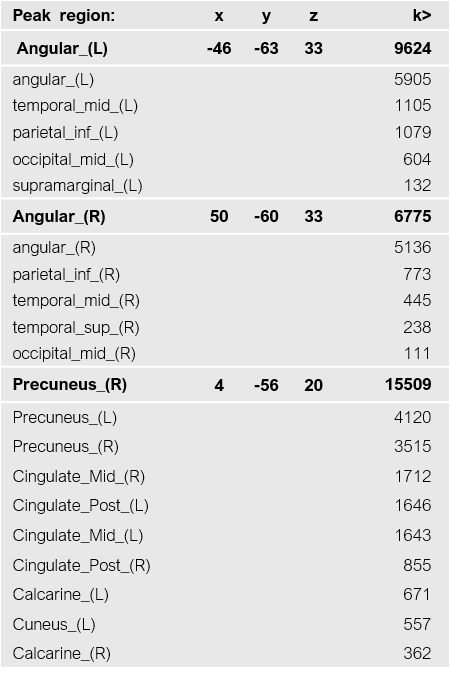

**Table S3.** Brain regions contributing to CAP1_DN-DM_. Peak MNI coordinates at z > 1.7 (*p<*.05) across all participants are reported for all regions*.* The names of the regions follow the Automated Anatomical Labelling Atlas 3 (AAL).

.

**CAP2 _DN-MT_**

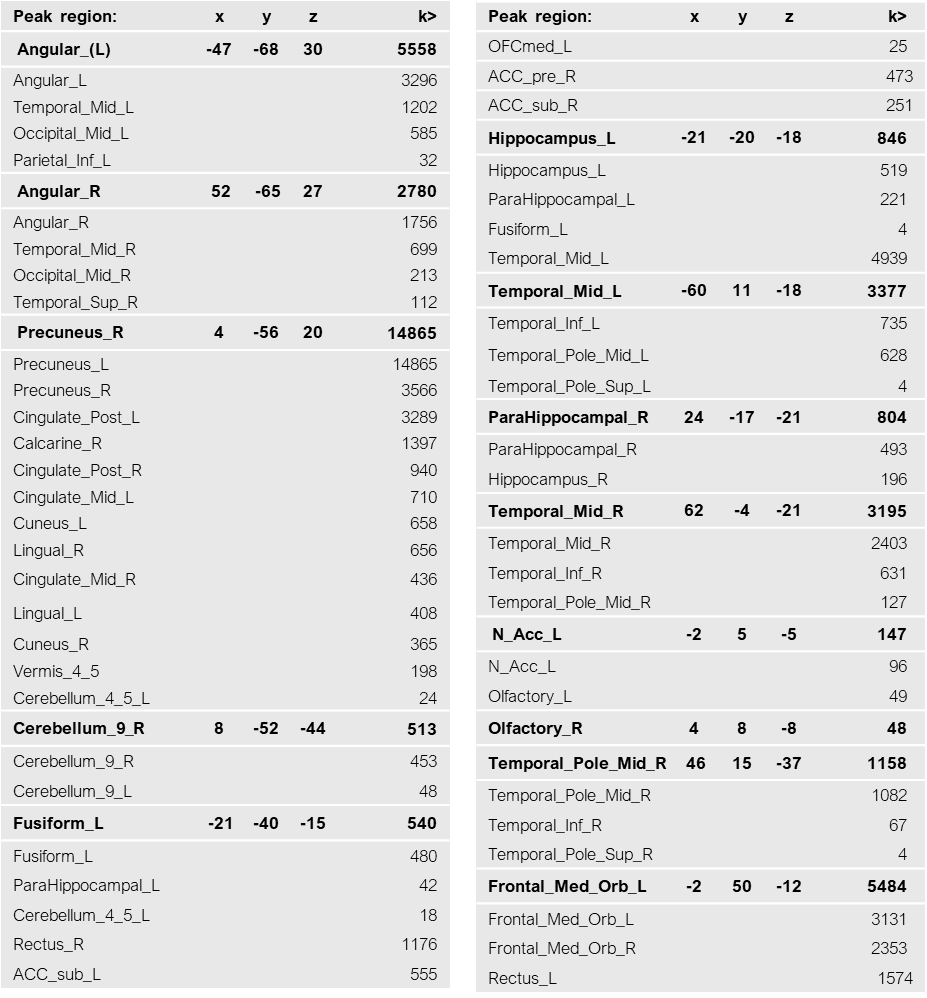

**Table S4.** Brain regions contributing to CAP_DN-MT_. Peak MNI coordinates at z > 1.7 (*p<*.05) across all participants are reported for all regions*.* The names of the regions follow the Automated Anatomical Labelling Atlas 3 (AAL).

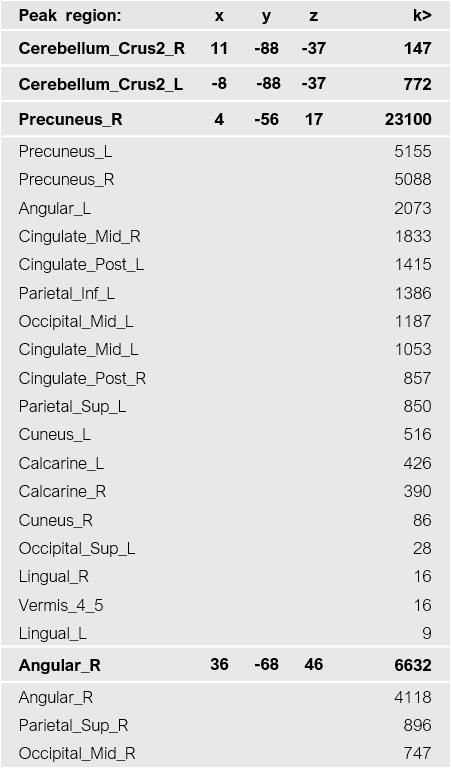

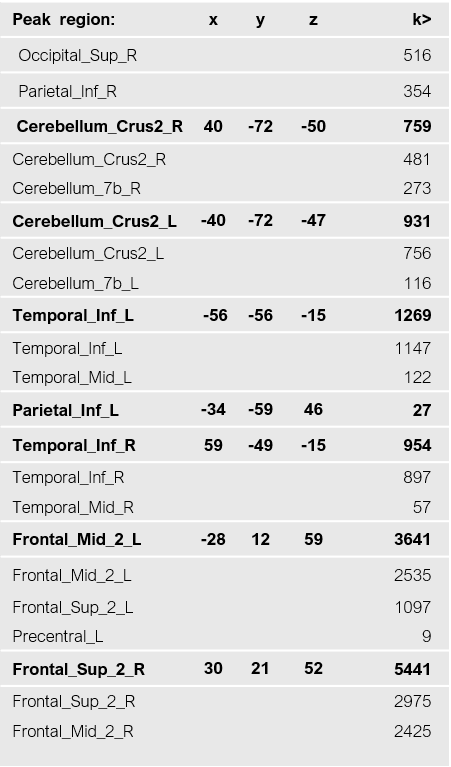

**CAP3_FPN_**

**Table S5.** Brain regions contributing to CAP_FPN_. Peak MNI coordinates at z > 1.7 (*p<*.05) across all participants are reported for all regions*.* The names of the regions follow the Automated Anatomical Labelling Atlas 3 (AAL).

**CAP4_VIS_**

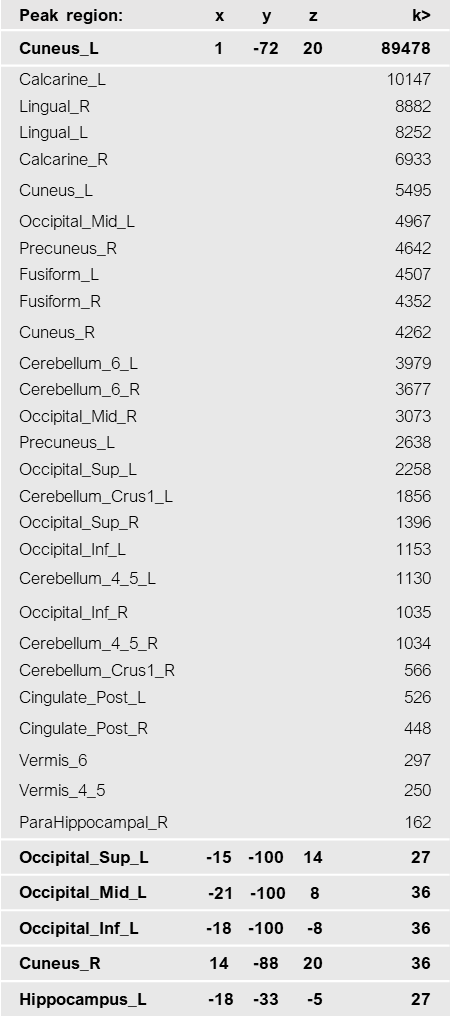

**Table S6.** Brain regions contributing to CAP4_VIS_. Peak MNI coordinates at z > 1.7 (*p<*.05) across all participants are reported for all regions*.* The names of the regions follow the Automated Anatomical Labelling Atlas 3 (AAL).

**CAP5_SN_**

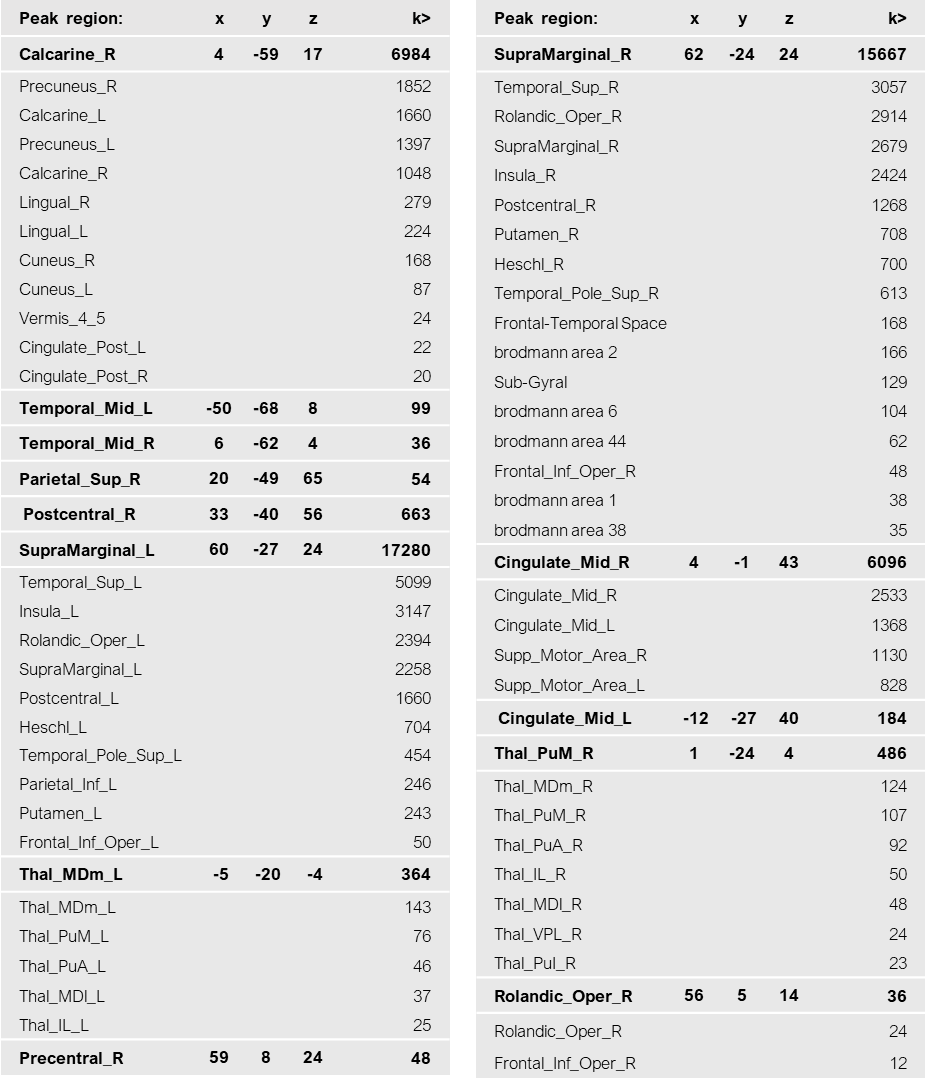

**Table S7.** Brain regions contributing to CAP5_SN_. Peak MNI coordinates at z > 1.7 (*p<*.05) across all participants are reported for all regions*.* The names of the regions follow the Automated Anatomical Labelling Atlas 3 (AAL).
